# KoVariome: Korean National Standard Reference Variome database of whole genomes with comprehensive SNV, indel, CNV, and SV analyses

**DOI:** 10.1101/187096

**Authors:** Jungeun Kim, Jessica A. Weber, Sungwoong Jho, Jinho Jang, JeHoon Jun, Yun Sung Cho, Hak-Min Kim, Hyunho Kim, Yumi Kim, OkSung Chung, Chang Geun Kim, HyeJin Lee, Byung Chul Kim, Kyudong Han, InSong Koh, Kyun Shik Chae, Semin Lee, Jeremy S. Edwards, Jong Bhak

**Author notes:** Contributed equally (Jungeun Kim, Jessica A. Weber, Sungwoong Jho). Corresponding authors (JB), (JE).

## Abstract

High-coverage whole-genome sequencing data of a single ethnicity can provide a useful catalogue of population-specific genetic variations. Herein, we report a comprehensive analysis of the Korean population, and present the Korean National Standard Reference Variome (KoVariome). As a part of the Korean Personal Genome Project (KPGP), we constructed the KoVariome database using 5.5 terabases of whole genome sequence data from 50 healthy Korean individuals with an average coverage depth of 31×. In total, KoVariome includes 12.7M single-nucleotide variants (SNVs), 1.7M short insertions and deletions (indels), 4K structural variations (SVs), and 3.6K copy number variations (CNVs). Among them, 2.4M (19%) SNVs and 0.4M (24%) indels were identified as novel. We also discovered selective enrichment of 3.8M SNVs and 0.5M indels in Korean individuals, which were used to filter out 1,271 coding-SNVs not originally removed from the 1,000 Genomes Project data when prioritizing disease-causing variants. CNV analyses revealed gene losses related to bone mineral densities and duplicated genes involved in brain development and fat reduction. Finally, KoVariome health records were used to identify novel disease-causing variants in the Korean population, demonstrating the value of high-quality ethnic variation databases for the accurate interpretation of individual genomes and the precise characterization of genetic variations.

## Introduction

The human reference genome^1^ was a milestone of scientific achievement and provides the foundation for biomedical research and personalized healthcare^2^. The completion of the human genome marked the beginning of our concerted efforts to understand and catalogue genetic variation across human populations. The International HapMap project resolved human haplotypes into more than one million common single nucleotide polymorphisms (SNPs) in an effort to catalogue genetic variations associated with diseases^3^. Subsequently, other large-scale genomic studies have identified 360M copy number variations (CNVs)^4^ and 6.4M small insertions and deletions (indels)^5^. These efforts laid the groundwork for approximately 1,800 genome-wide association (GWA) studies that investigated the genetic basis of complex diseases such as diabetes, cancer, and heart disease^6^. While these GWA studies have identified a wide range of disease-associated alleles that can be used as diagnostic tools^7^, the majority of the findings are associated with low disease risks and have led to a renewed focus on the detection of rare variants that are more predictive of disease^8^.

To identify pathogenic rare variants in GWA studies, disease cohorts are compared to population-scale variomes generated from healthy controls to remove common and low frequency variants in diverse human ethnic groups^9,10^. As a result, numerous population genomic studies have been performed to characterize ethnicity-relevant variations. One of the largest of such efforts, the 1,000 Genomes Project (1000GP), reported a total of 88M genetic variants, including SNPs, indels, and structural variations (SVs) from 2,504 healthy individuals^11^, and resolved population stratification by sampling 26 populations across five continents; Africa (AFR), East Asia (EAS), Europe (EUR), South Asia (SAS), and Americas (AMR). More recently, the Exome Aggregation Consortium (ExAC) released ten million human genetic variants from 60,706 individuals with a resolution of one exonic variant for every eight base-pairs^12^. Analysis of high coverage sequencing data (more than 30x) from 10,000 individuals showed that each newly analyzed genome added roughly 0.7MB of new sequences to the human reference genome and contributed an average of 8,579 new SNVs to the existing human variation data set ^13^. Large-scale variome studies, such as those previous discussed, have significantly increased our understanding of variation in the human population, however, the population composition is still broadly biased towards Europeans (54.97% in ExAC^12^ and 78.55% in Telenti *et al.*^13^). Consequently, many groups have initiated small variome studies of more targeted populations, i.e. the Malays^14^, Dutch (GoNL)^15^, Danish^16^, Japanese (1KGPN)^17^, Finland, and United Kingdom^18^. The large number of population-specific variations discovered in these studies highlights the importance of single population variomes in creating comprehensive databases of population heterogeneity and stratification.

SVs are also an important type of genomic variation in the human population that contribute significantly to genomic diversity^19^. SVs include large insertions (INSs), deletions (DELs), inversions (INVs), translocations, and CNVs^20^. Unlike SNVs and small indels, however, the identification of SVs remains challenging largely because of genome complexities and the limitations of short-read sequencing technologies^21^. Current efforts to resolve SVs reported several population-scale SVs^16,19^ and CNVs^17,22^ from whole genome sequencing (WGS) data, and these analyses characterized population-specific traits such as amylase gene duplication in high-starch diet populations^17,23^ and revealed associations for specific diseases such as hemophilia A^24^, hunter syndrome^25^, autism^26^, schizophrenia^27^, and Crohn’s disease^28^. Nevertheless, SVs identified in healthy individuals also contain a substantial number of individual- and population-specific SVs with no disease association. Taken together, these results have demonstrated the importance of constructing population-specific SV and CNV profiles for the precise characterization of disease association and identifying diagnostic markers for personalized medicine.

The Korean population is regarded as a relatively homogeneous ethnic group in East Asia^29^, from which a relatively small set of samples can produce a high-coverage population variome. Since the first Korean whole genome sequences were reported in 2009^30^, further variome studies in the Korean population have been conducted in the last decade using low-cost next generation sequencing (NGS) technologies^31-36^. Two exonic variomes of more than 1,000 Koreans were reported, though sampling was focused on disease cohorts containing patients with type II diabetes mellitus, hemophilia, cancer, and other rare diseases ^35,36^. Consequently, these studies are not suitable for parsing benign, demographic variants from disease variants. As the Korean center of the Personal Genome Project (PGP)^17^, the Korean Personal Genome Project (KPGP or PGP-Korea) was initiated in 2006 by the Korean Bioinformation Center (KOBIC) to resolve ethnicity-relevant variation in Korea by providing a comprehensive genomic, phenomic, and enviromic dataset accessible to researchers across the world. In 2009, KPGP published the first Korean genome with NGS data^30^ and the number of complete genomes has increased to 60 genomes as of 2016. This population was used to construct the first Korean Reference genome standard (KOREF)^37^, which was registered as a standard reference data for ethnic Korean genome sequence by evaluating its traceability, uncertainty, and consistency in the beginning of 2017.

To characterize the genomic variations across the Korean population, we selected and analyzed WGS data from 50 unrelated, healthy Korean individuals in KPGP cohorts with associated clinical diagnoses and family histories related to major diseases. In this report, we describe the general features of KoVariome and characterize all four types of genomic variations, which include 12.7M SNVs, 1.7M indels, 4K SVs, and 3.6K CNVs. This comprehensive database of genomic variations and corresponding metadata will be a valuable resource to the genomic community as researchers search for the genetic basis of disease.

## Results and Discussion

### Construction of the Korean standard Variome: KoVariome

Since 2010, the Korean variome data center, as a part of the KPGP, has been recruiting volunteers to generate WGS and whole exome sequencing (WES) data. The current KoVariome (version 20160815) has been constructed based on WGS data from 50 unrelated Korean individuals who responded to questionnaires detailing body characteristics, habits, allergies, family histories, and physical conditions related to 19 disease classes (Supplementary Table S1). A total of 5.5 TB of high-quality paired-end WGS data were generated, containing an average of 31× coverage per individual (Table 1 and Supplementary Table S2). WGS data from each individual covered 95% of the human reference genome (hg19) on average. From these data, we identified approximately 3.8M SNVs (ranged 3.7-3.9M) and 0.5M indels (0.4-0.7M) per Korean individual (Table 1 and Supplementary Fig. S1A). The hetero-to-homozygosity ratio of the autosomal SNVs was 1.49, which is consistent with previously reported data^38^. The length distributions of the indel loci were symmetric, with the majority of indel sizes shorter than six bases (94.8% for insertions, 97.8% for deletions) (Supplementary Fig. S1B). We identified approximately 20,097 (0.53%) SNVs and 258 (0.05%) indels in the coding regions including 10,394 (0.22%) non-synonymous changes per individual (Table 1).

**Table 1.** Statistics of KoVariome.

|  |  |
| --- | --- |
| Sample information for KoVariome |  |
| No. of samples (Male/Female) | 50 (31/19) |
| Total NGS yield | 5.5 tera bases |
| Average sequenced depth | 31x |
| Average mapped read rates | 95% |
| SNVs |  |
| Total No. of SNVs | 12,735,004 |
| No. of known variants in 1000GP <sup>a</sup> | 8,967,464 |
| No. of known variants in dbSNP <sup>b</sup> | 10,286,599 |
| Average No. of SNV per sample | 3,813,311 |
| Average No. of Coding SNVs <sup>c</sup> | 20,097 |
| Average No. of non-synonymous SNVs <sup>c</sup> | 10,394 |
| Average No. of SNVs with high effects <sup>c</sup> | 287 |
| Indels |  |
| Total No. of indels | 1,743,117 |
| No. of known variants in 1000GP <sup>a</sup> | 848, 471 |
| No. of known variants in dbSNP <sup>b</sup> | 1,307,000 |
| Average No. of indel per sample | 503,553 |
| Average No. of Coding indels <sup>c</sup> | 258 |
| Average No. of LOF indels <sup>c</sup> | 157 |

3 Variants deposited in <sup>a</sup> 1000GP and <sup>b</sup> the dbSNP (ver. 146). <sup>c</sup> predicted with SNPEff

Novel KoVariome SNVs were counted by adding individual samples one by one (Fig. 1A), and the number of novel SNVs logarithmically decreased and became depleted after the 9^th^ donor. In total, we observed 59K novel SNVs, including 1.2K (2.03%) coding-SNVs, per individual. To assess the relatedness of the KoVariome individuals, we compared the pairwise genetic distance of KoVariome with those of family data (Fig. 1B). WGS data from thirty families were downloaded from the KPGP database, which included two monozygotic twins, 14 parent-children pairs, seven siblings, five grandparents-grandchildren, six uncles-nephews, and three cousins. We analyzed familial SNVs using the same method as in KoVariome and also compared genetic distances between the two groups (see Methods). The genetic distance among KoVariome individuals was higher (pi=8.8e-4) than those found in the familial data, such as monozygotic twins (4.8e-4), siblings (6.7e-4), parent-child (6.8e-4), uncle-nephew (7.7e-4) and grandparents-grandchild (7.8e-4), and cousins (8.2e-4); verifying that no genetic bias was present in the sample collection stage. In accordance with previous reports, the multidimensional scaling (MDS) of the variants among Korean, Chinese, and Japanese individuals showed a clear separation of the three populations (Fig. S2) despite the geographical and historical associations between these groups^35,37^. These analyses reinforce the need for distinct KOREF and KoVariome reference resources to parse disease variants from demographic variants in this population.

**Figure 1.**
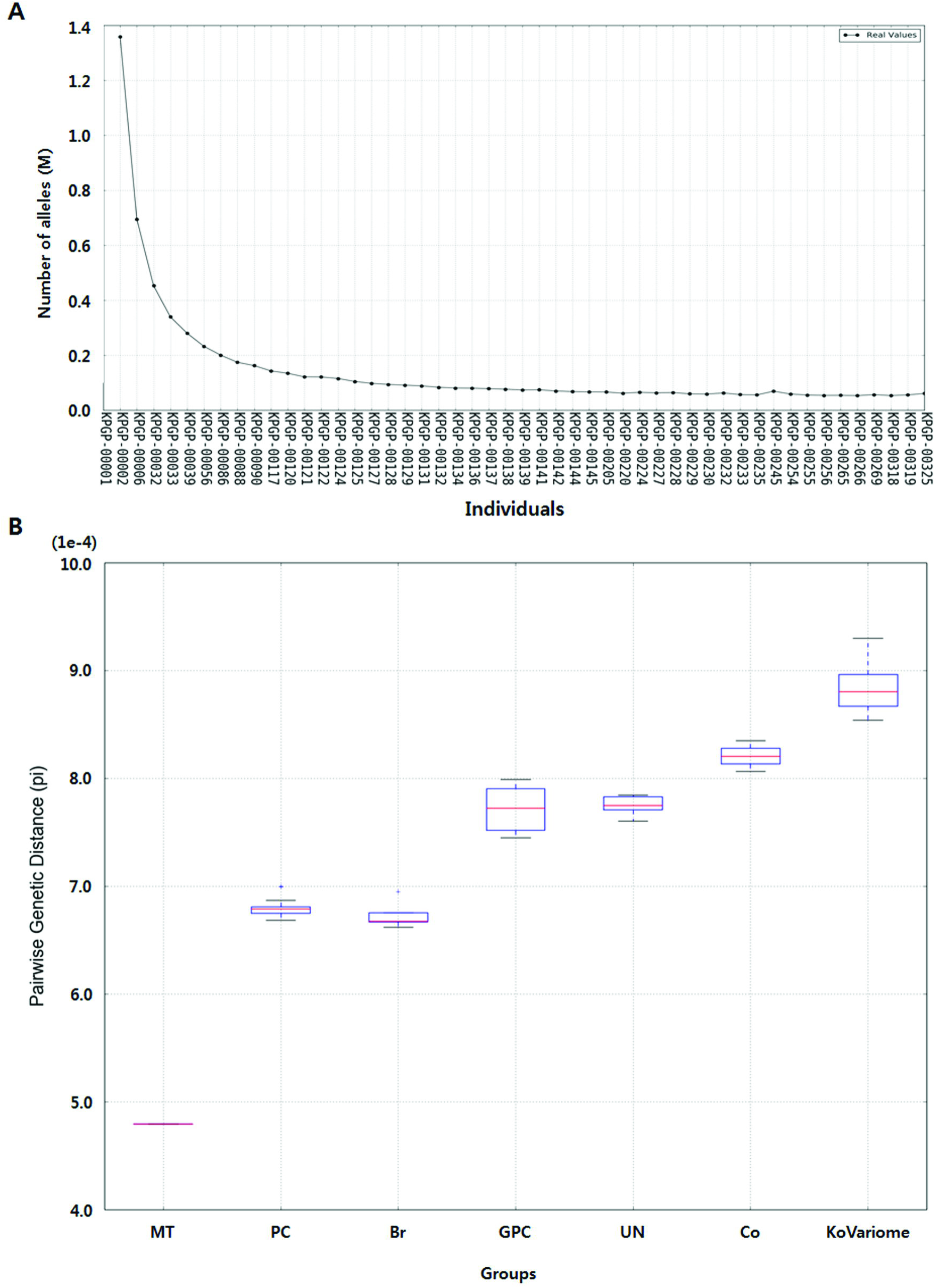
Status of KPGP variomes analyzed using 50 unrelated Korean individuals. A. Accumulation of novel SNV alleles. The number of novel SNV alleles were defined as newly identified nucleotides compared with previously constructed SNVs in KoVariome.
B. Genetic distance according to the familial relationships. Abbreviations: Monozygotic Twin (MT), Parent and child (PC), Brothers (Br), Grandparents vs. grand children (GPC), Uncle vs. Nephew (UN), and Cousins (Co)

### Accuracy test of SNVs and indels in KoVariome

We evaluated the accuracy of KoVariome SNV and indel predictions by comparing genotype results from the Axiom^TM^ Genome-ASI 1 Array with WGS data from 35 individuals. A total of 503,694 SNV positions were compared, from which we obtained an average of 0.9993 precision (ranged: 0.9984-0.9996) and 0.9980 recall (ranged: 0.9817-0.9994) (Supplementary Table S3). In addition, there was a 99.65% (ranged: 98.62-99.87%) concordance of the SNVs called by the WGS and Axiom array calls. Compared to similar variome studies, this genotype accuracy was slightly lower than the high-depth trio data in the Danish population study (99.8%)^16^ but higher than that of the Dutch population SNVs (99.4-99.5%) analyzed with intermediate depths^39^. The accuracy of the SNV calls was analyzed across the genome, and a total of 499,889 (99.24%) SNVs showed a genotype concordance higher than 0.99, while 0.4% of SNVs showed the genotype accuracy less than 0.95 (Supplementary Table S4). Similar levels of genotype concordances were observed in the repetitive regions of the genome (99.56% of SNVs with the genotype correspondence > 0.95, Supplementary Table S5), suggesting that SNV calling accuracy is not reduced in repetitive regions of the genome.

We also compared the accuracy of indel variant calls with the 1,981 indel markers on the Axiom^TM^ Genome-ASI 1 Array. A genotype comparison showed an average accuracy of 98.49% for indels, which was slightly lower than those observed in SNVs (Supplementary Table S3), and comparable to the false positive (FP) rate for indels that was reported in the Danish data^16^. In terms of genomic loci, 1,343 (91.11%) indels showed perfect genotype concordance with array data and 1,446 (98.10%) indels had an accuracy higher than 90% (Supplementary Fig. S3).

### Genome-wide features of KoVariome

By merging the variants of 50 unrelated Korean individuals, we identified 12.7M SNVs and 1.7M small indels shorter than 100bp (Table 1); approximately 1.5 times the number of SNVs previously reported from preliminary KPGP data (0.8M)^33^. Both types of variants were primarily distributed in the non-coding regions (about 98%), including intergenic and intron regions (Supplementary Table S6). Approximately 10.3M (81.10%) SNVs and 1.3M (76.47%) indels were present in dbSNP (ver. 146); while 2.4M SNVs and 0.4M indels were novel (Table 1). A total of 9M (70.42%) SNVs and 0.8M (48.68%) indels were found in the 1000GP variome (Supplementary Table S6); and based on allele frequencies, 4.6M (51.03%) and 4.4M (48.82%) of these SNVs were classified into the categories ‘1000GP common’ and ‘1000GP low frequency’, respectively (Fig. 2A). Most notably, 13,584 (0.15%) KoVariome SNVs were rarely observed in the 1000GP continental groups with a MAF < 0.1%. A similar distribution was observed with the indels, where 64.2% and 35.8% of the KoVariome indels were classified into the ‘1000GP common’ (0.5M) and ‘1000GP low frequency’ (0.3M) classes, respectively. Only ten indels were classified into the ‘1000GP rare’ category. Almost all of the variants in the ‘1000GP common’ category were also frequently observed in KoVariome, representing 4.5M (98.33%) SNVs and 0.5M (93.37%) indels in this class (Fig. 2A and Supplementary Table S6). Surprisingly, however, roughly half of the variants in ‘1000GP low frequency’ were classified as ‘frequent in KoVariome’.

**Figure 2.**
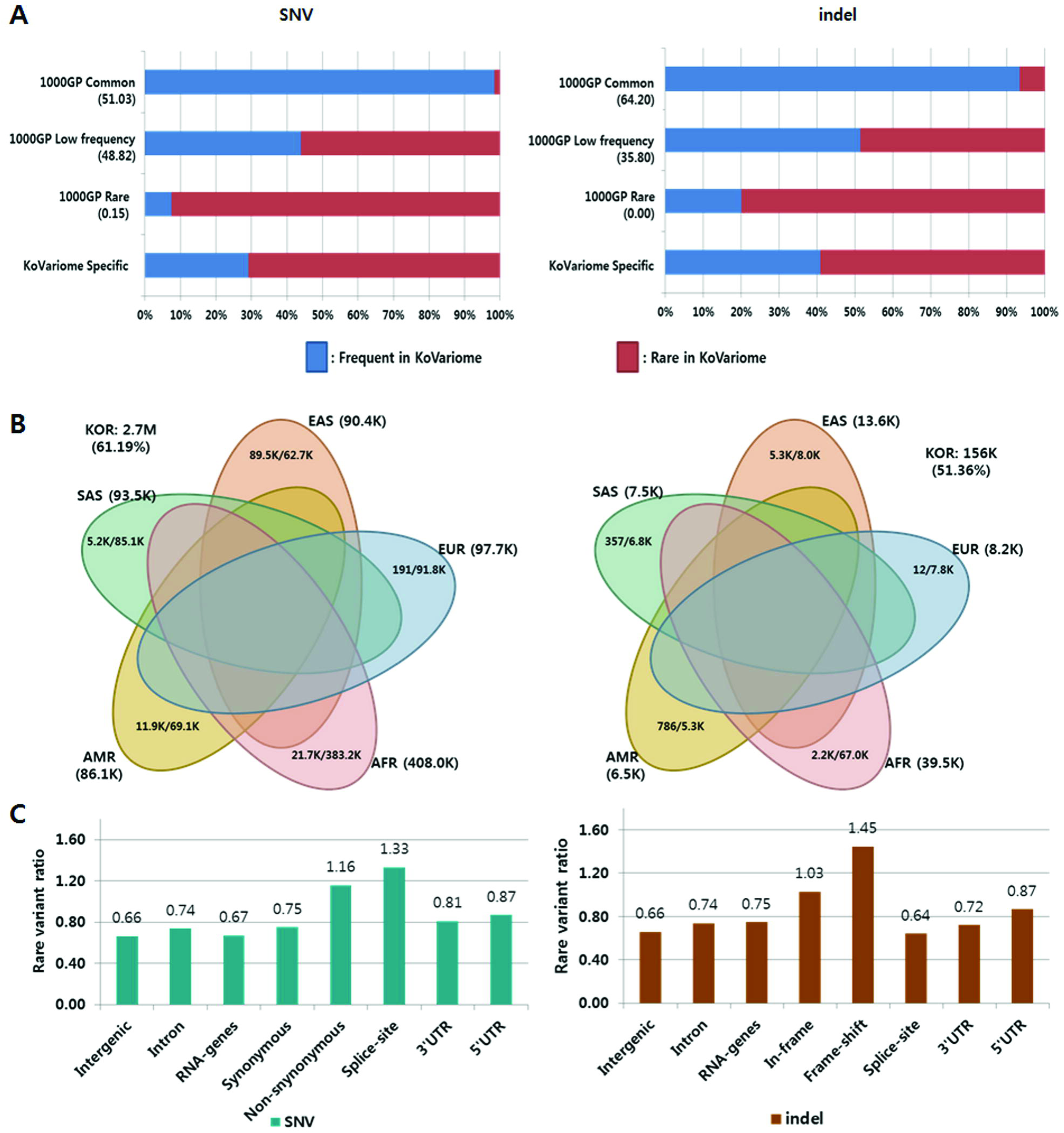
Genetic features of KoVariome. A. Two dimensional classification of KoVariome. SNVs and indels observed in 1000GP data were classified based on the minor allele frequencies (MAF); ‘1000GP Common’: MAF >=5% in all five continents, ‘1000GP Low frequency’: MAF >= 0.1% in any continent, and ‘1000GP Rare’; MAF < 0.1% in all five continents. The five continental populations included African (AFR), European (EUR), Native American (AMR), South Asian (SAS), and East Asian (EAS). The second group was classified by the number of variants in KoVariome; ‘Frequent in KoVariome’ (>= 3) and ‘Rare in KoVariome’ (< 3). B. The Venn diagrams represent the number of variants enriched in specific continents for both SNVs (left) and indels (right). The enrichment was analyzed by Fisher’s exact test based on odds ratio > 3 and p-value < 0.01. The total numbers of enriched variants in the Korean (KOR) population are denoted in the white space of the Venn diagram. The numbers next to the continental population abbreviations represent the total number of enriched variants in that 1000GP continental group. The numbers within each ellipse denote the number of variants enriched both in KOR and a specific continent (left) and the number of variants enriched exclusively in the represented continent (right). C. Rare variant ratios (RVRs) observed in each genomic region. RVRs were calculated by dividing the number rare variants by the number of frequent variants in KoVariome.

Next, we compared the allele frequencies in five the continental 1000GP groups to KoVariome. In total, we observed 3.4M (77.19%) SNVs and 0.2M (74.21%) indels that were statistically enriched in at least one of the continental groups or the Korean population (Fig. 2B), suggesting a population stratification. To further explore the population stratification, we identified the variants uniquely enriched in each continental group, and the enriched variants that were in common between the continental groups. In total, nearly three million (2.7M) SNVs and 156K indels were frequently found in the Korean population. Among them, 2.5M (95.20%) SNVs and 143K (94.47%) indels showed Korean specific enrichments, while the other enriched variants were shared by other continents (Figure 2B). Among the five continental groups, as expected, EAS shared the largest number of enriched variants (89.5K SNPs and 5.3K indels) with the Korean population ^40^.

### Interpretation of the KoVariome-specific variants

Characterizing ethnicity specific variants is necessary to understand the demographic differences between populations and can be used to filter out low frequency clustered variants in a specific group. In KoVariome, there were 3.8M SNVs and 0.9M indels not observed in the 1000GP variome (Supplementary Table S6). Among them, 1.1M (29.16%) SNVs and 0.4M (40.88%) indels were classified as ‘frequent in KoVariome’ (Fig. 2A). Of the 15,279 non-synonymous SNVs and 480 frame-shift indels specific to KoVariome, 11,746 (76.88%) and 397 (82.71%) were rare in KoVariome (n < 3), respectively; whereas 3,533 SNVs were frequently observed (occurring at least three times) in KoVariome but not observed in the 1000GP variome.

To identify the clinical relevance of these variants, we compared the genomic loci of these SNVs against the ClinVar database and identified six pathogenic or likely pathogenic loci with associated disease information (Table 2). Two pathogenic SNVs (rs386834119 and rs1136743) were autosomal recessive (AR) diseases, and therefore, no phenotypes were expected since all of the KoVariome SNVs were heterozygotes in the KPGP. We observed a high allele frequency (three males and two females) of the cancer-associated SNV (rs200564819) in *RAD51*, which is known to increase the risk of developing ovarian and breast cancers^41^. While the inheritance type is not available in the Online Mendelian Inheritance in Man (OMIM), we speculate that it is autosomal dominant (AD) with incomplete penetrance since four out of 14 male *RARD51*-deficient carriers (heterozygous) were diagnosed with colorectal cancers. However, none of the donors with this SNV have been diagnosed cancer and have no familial cancer history. We also observed a splicing-donor (‘GU’) candidate five nucleotides downstream of this SNV, although further confirmation is required. This site may induce the null effect of rs200564819 by the creation of new splice-sites according to the guidelines from the American College of Medical Genetics and Genomics (ACMC)^42^. In addition, we observed two pathogenic missense SNVs (rs121912678 and rs20016664) associated with fibrodysplasia ossificans progressive (FOP) and Van der Woude syndrome (VWS), respectively (Table 2). A chr2:g158630626C>G SNV was rarely observed in the ExAc database (MAF=0.0002) and another variant (C>T) at this position revealed a pathogenic effect for FOP disease by changing R206H in the *activin receptor type I* (*ACVR1*) gene^43^. While the pathogenicity of R206P in *ACVR1* due to a C>G mutation is not yet known, we suggest that it is likely benign because of the high MAF (0.14) of this allele without any FOP phenotypes, skeletal malformation, or progressive extraskeletal ossification recorded in the KPGP survey. Furthermore, the 400^th^ amino acid of the *interferon regulatory factor 6* (*IRF6*) gene is known to be a hot spot of VWS, orofacial clefting disorders. Two pathogenic residues, R400W^44^ and R400Q^45^, were reported for VWS; however, the pathogenicity of R400P arisen by chr1:209961970C>G is not yet confirmed. A total of 14 heterozygous SNVs had no phenotype for VWS symptom, despite the AD inheritance pattern of this disease; and consequently, the R400P substitution also seems to be benign. Taken together, the KoVariome-specific frequent variants demonstrate the importance of using population-scale health data to identify pathogenic loci in specific diseases, and for the accurate identification of benign variants that are not annotated because of population stratification.

**Table 2.**
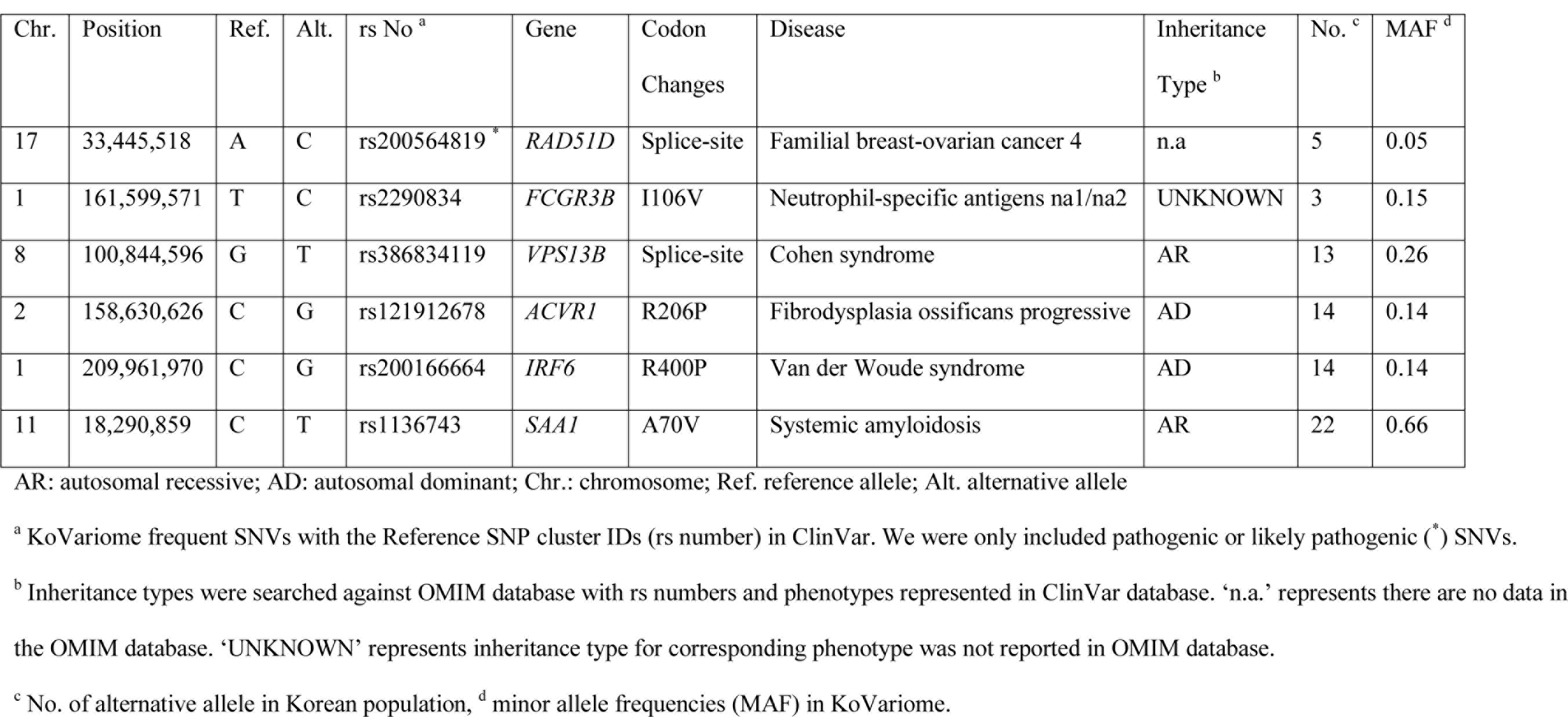
ClinVar annotation of the KoVariome frequent SNVs.

### Functional impact of rare variants

We investigated the proportion of the SNVs in four SNV classes (1000GP Common, 1000GP Low Frequency, 1000GP Rare, KoVariome Specific; Supplementary Fig. S4). Our analyses showed that a higher portion of the coding SNVs were enriched in the ‘1000GP rare’ class, while the SNVs in the non-coding regions were similarly distributed in all other variant classes. The portion of non-synonymous SNVs in the ‘1000GP rare’ class was more than twice what was observed in the other classes. It is possible that these patterns are associated with purifying selection to rapidly remove deleterious alleles in the population^46^, though it was not possible to identify this pattern in frame-shift indels because of the small number of variants (981) in this class. To analyze the tendencies of purifying selection in KoVariome, we defined rare variant ratios (RVRs) as the number of SNVs in the ‘rare in KoVariome’ class divided by the number of SNVs in the ‘frequent in KoVariome’ class. We then compared RVRs across genomic regions (Fig. 2C). In both SNVs and indels, RVRs in the intergenic region were lowest (0.66), while similar levels of RVRs were observed in other non-coding regions (0.66-0.87). Under the assumption that mutations occur randomly throughout the genome, lower rates of RVR in non-coding regions suggest neutral selection with no or weak selection pressures in the population. Conversely, the highest RVR of frame-shift indels (1.45) suggests there was some purifying selection against these variants in the Korean population. Furthermore, about twice as many RVRs were observed in the non-synonymous (1.16) and splice-site (1.33) SNVs compared to intergenic regions. Although SNVs in the coding region can be deleterious to protein function, selection pressure on the non-synonymous and splice-site SNVs seem to be slightly lower than that of the frame-shift indels.

### Interpretation of disease-causing variants among Korean individuals

Rare SNVs in an individual genome are more likely to be pathogenic than common variants. Because genetic variants are known to be geographically clustered, characterizing population stratification is a critical first step to identifying disease-causing variants^47^. With this concept, we examined rare SNVs in each individual after filtering out common SNVs that were classified as ‘1000GP common’, ‘1000GP low frequency’, or ‘frequent’ in KoVariome. From an average of 3.8M SNVs per individual, 3.4M (88.70%) and 0.4M (9.39%) SNVs were filtered out using the 1000GP variome or KoVariome, respectively (Fig. 3A and Table 3). Overall, KoVariome allowed 1,231 (12.25%, median value) non-synonymous SNVs and 40 (24.01%) splice-site SNVs to be filtered out as common variants in the Korean population, which significantly improves the ability to pin-point disease causative variants.

**Figure 3.**
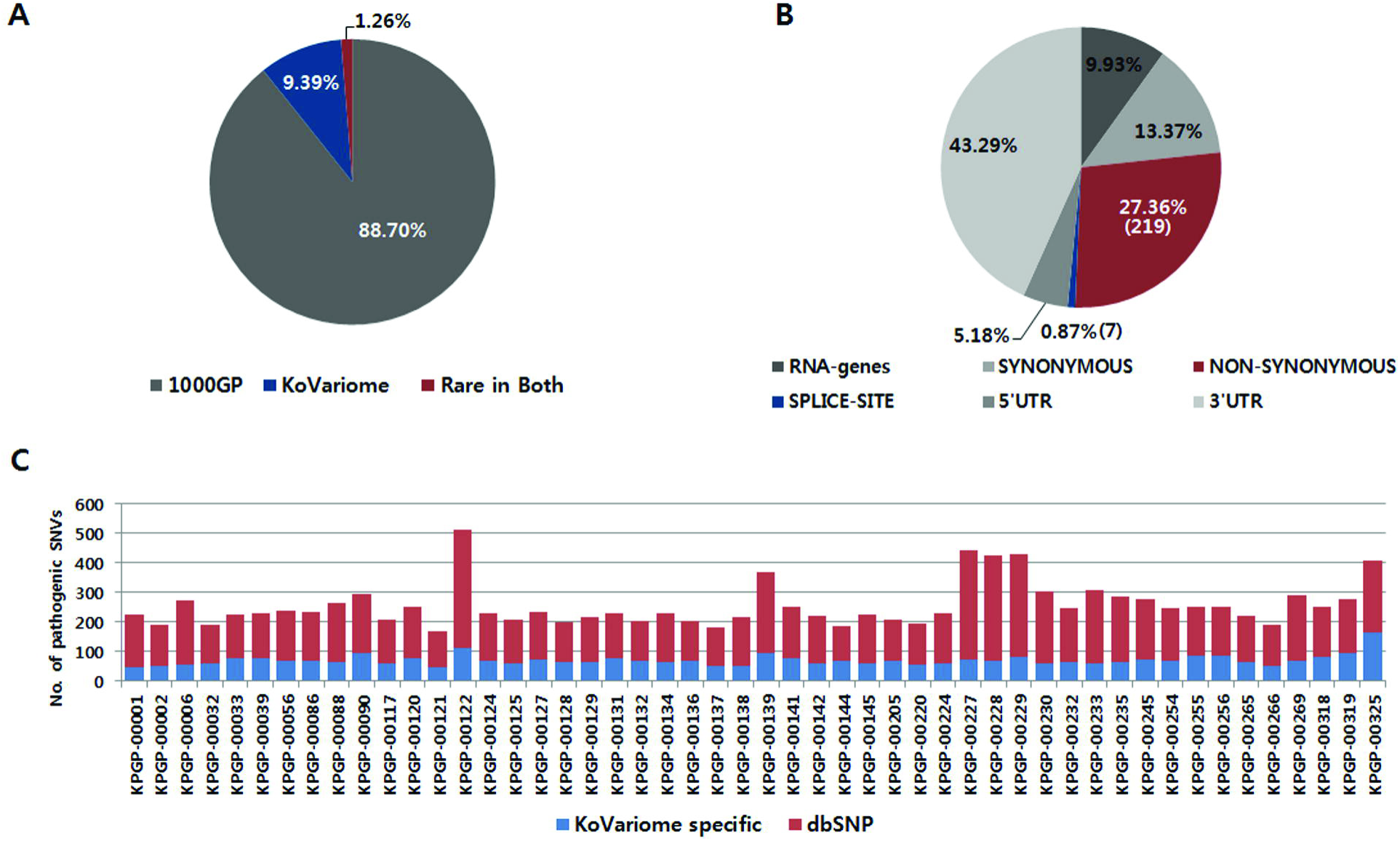
Individual variants describing functional effects. A. Classification of individual variants based on frequency in 1000GP and KoVariome. Gray represents the portion of individual variants classified in the ‘1000GP common’ and ‘1000GP Low frequency’. Blue represents the portion of the individual variants classified in the ‘Frequent in KoVariome’. Red represents rare variants in both 1000GP and KoVariome ‘Rare in Both’. B. Individual variants in the ‘Rare in Both’ were classified by gene coordinates. To more clearly represent the portion functionally important rare variants, 98% of the rare variants in the non-coding regions were not represented. C. Number of pathogenic variants for each individual. Red and blue bars represent the number of pathogenic variants previously reported in dbSNP and novel, respectively.

**Table 3.**
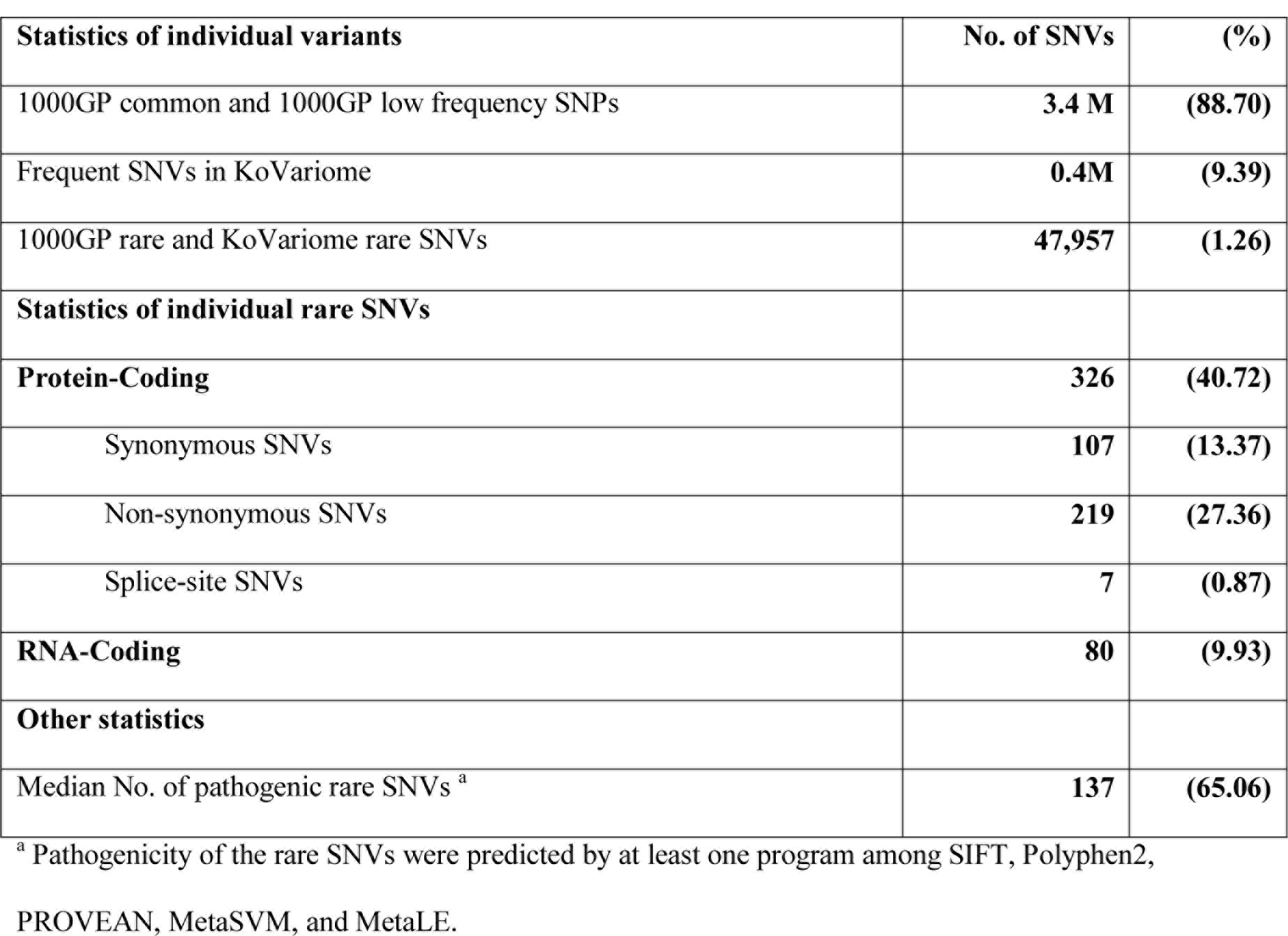
Statistics of individual SNVs.

| **Statistics of individual variants** | **No. of SNVs** | **(%)** |
| --- | --- | --- |
| 1000GP common and 1000GP low frequency SNPs | **3.4 M** | **(88.70)** |
| Frequent SNVs in KoVariome | **0.4M** | **(9.39)** |
| 1000GP rare and KoVariome rare SNVs | **47,957** | **(1.26)** |
| **Statistics of individual rare SNVs** |  |  |
| **Protein-Coding** | **326** | **(40.72)** |
| Synonymous SNVs | **107** | **(13.37)** |
| Non-synonymous SNVs | **219** | **(27.36)** |
| Splice-site SNVs | **7** | **(0.87)** |
| **RNA-Coding** | **80** | **(9.93)** |
| **Other statistics** |  |  |
| Median No. of pathogenic rare SNVs <sup>a</sup> | **137** | **(65.06)** |

<sup>a</sup> Pathogenicity of the rare SNVs were predicted by at least one program among SIFT, Polyphen2,
PROVEAN, MetaSVM, and MetaLE.

After filtering, Korean donors had a median of 47,957 (1.26%) rare SNVs, most of which (98.33%) were located in non-coding regions. Among these rare SNVs, we observed an average of 219 (67.17%) non-synonymous SNVs and seven (0.87 %) splice-site SNVs per individual (Fig. 3B and Table 2). On average, 166 (73.45%) of these SNVs were present in dbSNP (ver. 146), but not in the 1000GP variome (Fig. 3C). Of the 12,445 non-synonymous rare SNVs distributed in 50 Korean individuals, we identified 7,645 (61.43%) pathogenic or probably pathogenic SNVs predicted by at least one computational algorithm (see methods section, Table S7). In total, 38 (0.5%) pathogenic rare SNVs in KoVariome were homozygotes and the remaining (99.5%) were heterozygotes. In addition, 29 (58%) of the donors had no homozygous pathogenic rare SNVs. To obtain clinical information concerning these pathogenic rare-SNVs, we searched the genomic loci for these SNVs against the ClinVar database. A total of 127 of the rare SNVs were found in ClinVar, 53 of which showed clear clinical significance. Eight (6.39%) and thirteen (10.24%) were listed as benign and likely benign in ClinVar, respectively, and not fatal for a specific disease. Conversely, 29 (22.83%) and three (2.36%) were pathogenic and likely pathogenic, respectively (Table 4). These rare SNVs contribute to disease according to their inheritance patterns, and a manual investigation of the inheritance type using the OMIM database identified seven AD and 17 AR SNVs for specific loci; although we failed to identify the inheritance types for eight SNV loci (Table 4). All 17 of the AR SNVs were heterozygous in KoVariome, so it was not possible to assign phenotypes to these loci. Within the donor group with pathogenic rare AD SNVs, we searched for phenotypes or familial histories associated with target diseases in the questionnaire. We identified a familial history for type II diabetes mellitus associated with rs121918673 allele KPGP participants; however, one donor with the rs121918673 allele was nondiabetic and reported no family history of this disease Additionally, one donor was heterozygous for the rs121912749 allele, which has been associated with spherocytosis, and this donor reported associated symptoms but no anemia (Supplementary Table S1 and S7). However, it is clinically known that spherocytosis has heterogenetic symptoms ranging from asymptomatic to hemolytic anemia. These examples highlight the disease-relevant genetic information this resource can provide to patients, and emphasize the utility of KoVariome to the Korean population at large as WGS becomes a more routine component of healthcare.

**Table 4.** Known pathogenic rare variants associated with disease.

| Individual ID | rs No. | Genotype | Codon change | Inheritance type <sup>a</sup> | gene | ClinVarTraits |
| --- | --- | --- | --- | --- | --- | --- |
| KPGP-00001 | rs563607795 | A/G | L385P | n.a. | *SLC19A3* | Thiamine metabolism dysfunction syndrome |
| KPGP-00001 | rs199769221* | G/C | R116P | AD | *PRSS1* | Hereditary pancreatitis |
| KPGP-00032 | rs387907164 | T/C | C32R | AR | *KIAA1530* | UV-sensitive syndrome 3 |
| KPGP-00033 | rs119490107 | C/A | D234Y | UNKNOWN | *RAD54B* | Carcinoma of colon |
| KPGP-00039 | rs199476197 | A/C | H331P | AR | *CYP4V2* | Bietti crystalline corneoretinal dystrophy |
| KPGP-00088 | rs28940280 | G/A | D279N | AR | *CLN5* | Ceroid lipofuscinosis neuronal 5 |
| KPGP-00122 | rs587782989 | C/T | R464H | AD | *CCDC88C* | Spinocerebellar ataxia 40 |
| KPGP-00124 | rs142808899 | C/T | G303R | AR | *DHCR7* | Smith-Lemli-Opitz syndrome |
| KPGP-00127 | rs111033744 | A/G | Y100C | AR | *GALT* | Galactosemia |
| KPGP-00127 | rs137852972 | T/C | N88S | AD | *BSCL2* | Silver spastic paraplegia syndrome |
| KPGP-00129 | rs137853022 | C/T | R696Q | AR | *IKBKAP* | Familial dysautonomia |
| KPGP-00129 | rs386833823* | G/A | S238F | AR | *SLC7A7* | Lysinuric protein intolerance |
| KPGP-00131 | rs200088377 | G/A | P191L | n.a. | *IL17RD* | Delayed puberty |
| KPGP-00136 | rs121908099 | G/A | R405Q | AR | *CYP27A1* | Cholesterol storage disease |
| KPGP-00136 | rs750218942 | C/G | Splice-site | AR | *XPA* | Xeroderma pigmentosum |
| KPGP-00136 | rs727502791 | G/A | R158* | AD | *MFAP5* | Aortic aneurysm (familial thoracic 9) |
| KPGP-00136 | rs545215807 | G/A | G109S | AR | *ACADVL* | VLCAD deficiency |
| KPGP-00139 | rs387907033 | G/C | G401A | AR | *SYT14* | Spinocerebellar ataxia |
| KPGP-00139 | rs748486078 | G/A | S95L | UNKNOWN | *IL17F* | Candidiasis |
| KPGP-00144 | rs119480073 | C/T | R80I | AR | *LPIN1* | Myoglobinuria |
| KPGP-00144 | rs104895438 | G/A | A612T | AD | *NOD2* | Sarcoidosis |
| KPGP-00205 | rs121913050 | G/A | R153H | UNKNOWN | *ERCC4* | XFE progeroid syndrome |
| KPGP-00220 | rs121918673 | G/C | S439R | AD | *HNF1B* | Diabetes mellitus type 2 |
| KPGP-00266 | rs104894085 | G/A | Q258* | AR | *STAR* | Cholesterol monooxygenase deficiency |
| KPGP-00227 | rs121909569 | A/G | S148P | AD, AR | *SERPINC1* | Antithrombin III deficiency |
| KPGP-00228 | rs121434426 | G/A | Q356* | UNKNOWN | *FANCG* | Fanconi anemia |
| KPGP-00232 | rs121909385 | T/C | L623P | AR | *SLC12A3* | Familial hypokalemia hypomagnesemia |
| KPGP-00233 | rs672601312 | G/T | E127* | AR | *ISG15* | Immunodeficiency 38 with basal ganglia calcification |
| KPGP-00233 | rs749462358 | C/T | E924K | n.a. | *ASPM* | Not provided |
| KPGP-00245 | rs137854500 | C/T | D1289N | AR | *ABCA1* | Tangier disease |
| KPGP-00254 | rs201968272 | G/A | R237Q | AR | *DDX11* | Warsaw breakage syndrome |
| KPGP-00325 | rs121912749 | C/T | G130R | AD | *SLC4A1* | Spherocytosis type 4 |

Abbreviations: Chr. chromosome; Ref. reference allele; Alt. alternative allele; AD: autosomal dominant; AR: autosomal recessive
\* The clinical significance of SNV locus was defined as likely pathogenic in the ClinVar database.
<sup>a</sup> Inheritance type were searched against OMIM database with rs numbers and phenotypes in the ClinVar database. ‘n.a.’ represents there are no data in the OMIM database. ‘UNKNOWN’ represents inheritance type for corresponding phenotype was not reported in OMIM database.

### Structural variations in KoVariome

SVs are common across the human genome, though identifying and defining the impact of SVs is more difficult than SNVs or indels (<100bp). We predicted on average 6,534 individual SVs, including 450 INVs, 354 intra-chromosomal translocations (ITXs), 478 INSs, and 5,252 DELs using BreakDancer (BD) and Pindel programs (Supplementary Table S8). To identify SVs with clear break points, we removed 15-32% spurious SVs per individual (see Methods; Supplementary Fig. S5 and Table S8). After filtering, we obtained 40,179 non-redundant SVs; including 4,896 INVs, 2,131 ITXs, 12,171 INSs, and 20,981 DELs. Within the Korean donor group, individuals contained 3,294 SVs (median), 82.36% of which were DELs (Fig. 4A). The median length of individual SVs was 2.3Kb for INVs, 5.8Kb for ITXs, 1.3Kb for INSs, and 342bp for DELs (Fig. 4B). A high proportion of SVs were specific to an individual genome (Fig. 4C), consistent with findings from the 1KJPN^17^. The portion of individual-specific SVs was greatest for INSs (92.51%), followed by INVs (88.87%), ITXs (68.93%), and DELs (47.82%) (Table S8). A substantial proportion of SVs (98.5% INSs and 61% DELs) were novel and were not previously deposited in the Database of Genomic Variants (DGV). Overall, the non-redundant combined SVs ranged in size up to 10M and all classes were enriched in the 1-2Kb size range (Fig. 4D, Supplementary Fig. S6).

**Figure 4.**
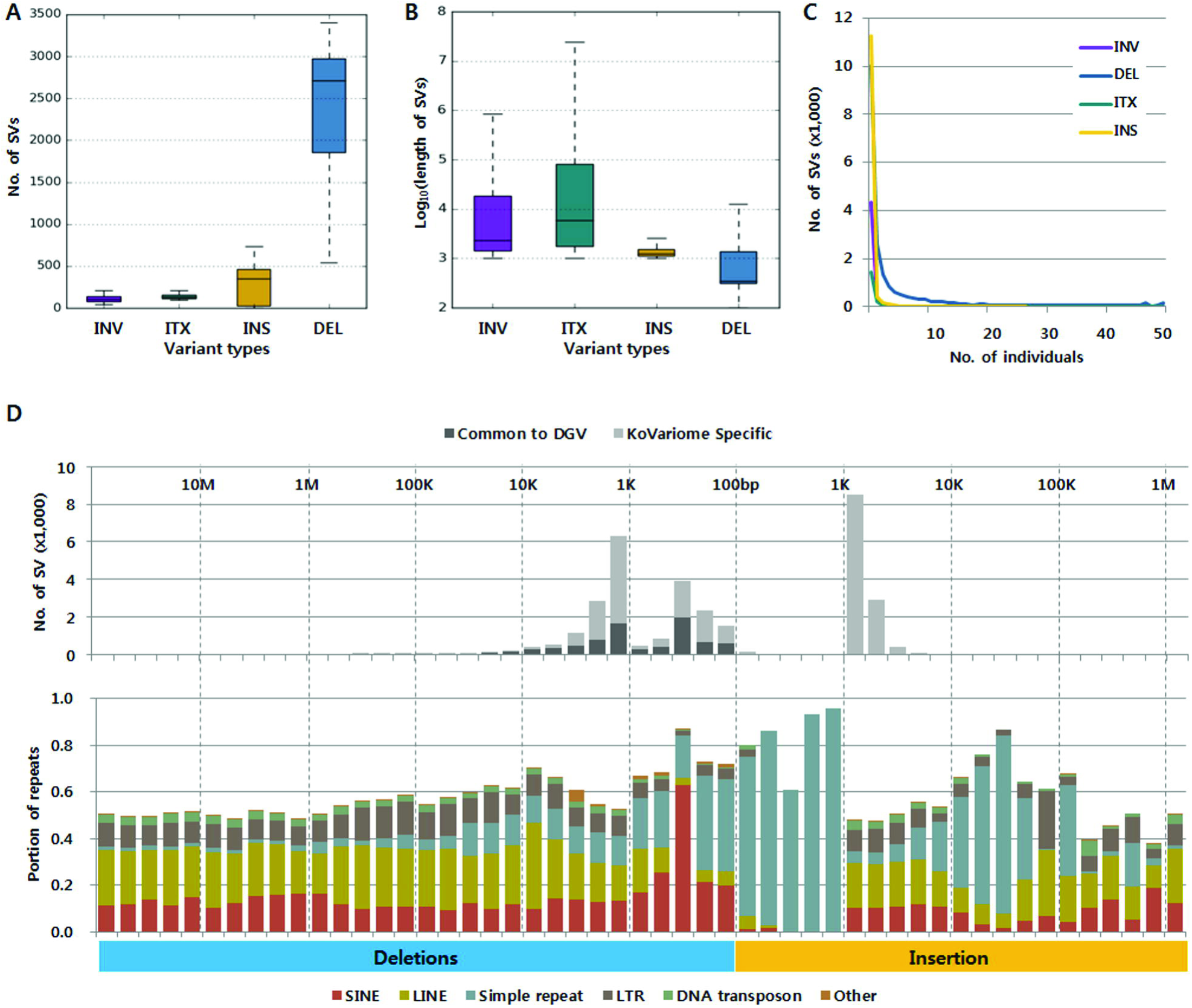
Properties of structural variants discovered in KoVariome. A. The boxplot represents the number of variants per Korean individual by variant type (n=50). The lower and upper hinges of the boxes correspond to the 25^th^ and 75^th^ percentiles and the whiskers represent the 1.5x inter-quartile range (IQR) extending from the hinges. Abbreviations of the variants: inversions (INV), intra-chromosomal translocation (ITX), insertions (INS), and deletions (DEL). B. Length of the variants present in the individual genome. See variant types and boxplot definition in A. C. Frequency of variants in KoVariome. D. The upper graph represents the number of SVs identified at specific length ranges. The KoVariome specific variants were defined by comparing SVs in the Database of Genomic Variants (DGV) with 70% reciprocal overlap. The lower graph represents the portion of repeats distributed in the variants. Repeat classes were defined by the repeat annotations provided in the UCSC Genome bioinformatics. Simple repeats contained both microsatellites and low complexity (e.g., AT-rich). Abbreviations of repeats: short interspersed element (SINE), long interspersed element (LINE), and long terminal repeat (LTR).

Finally, we analyzed the SVs to determine whether they were enriched for repetitive elements. Within the SVs, we cataloged repeat types and searched for Korean-specific enrichments compared to those present in other populations. Among the SVs, we found that 13% contained short interspersed elements (SINEs), 20% contained long interspersed elements (LINEs), 3.4% contained DNA transposons, and 8.6% contained long terminal repeats (LTRs). The majority of SINEs were observed in DELs of 200-300bp, which is consistent with *de novo* assembled SVs^16^ and the predicted SVs^15^. These results suggest that SVs are enriched for SINEs in the 1-4Kb INVs, and LINEs in the 4-40Kb INVs (Supplementary Fig. S6A). Additionally, simple repeats were predominantly observed in INSs (Fig. 4D) and 3-5Kb ITXs (Supplementary Fig. S6B).

### Copy number variations in KoVariome

The high coverage WGS data used to construct KoVariome provides sufficient data to characterize CNVs in a single genome. The FREEC program^48^ predicted an average of 199 deletions and 336 duplications per genome (Supplementary Table S9). After filtering out spurious CNVs (Supplementary Fig. S7), 161.74 (81.46%) deletions and 296.72 (88.29%) duplications remained from the original calls. In total, we predicted 2,038 non-redundant deletions and 1,564 non-redundant duplications, and the unified CNVs were approximately 5Kb-100Kb in length (Fig. 5A). When compared to the DGV, we identified 3.6K known CNVs, including 1,169 (57.36%) deletions and 846 (54.09%) duplications. Repeat composition analyses of CNV regions revealed that deletions smaller than 5K and duplications smaller than 10K contained a 20-fold more simple repeats compared to their overall frequencies in the human genome. In addition, SINEs were 2-fold more frequent in the > 600Kb deletions. These associations differ from the repeat distributions in SVs. By examining the genes in the unified CNVs, 869 (46.47%) deletions and 1,105 (70.65%) duplications were found to contain at least one gene. In addition, only two deletions and three duplications were conserved in the 50 Korean individuals (Table 5). Interestingly, a long 2M genomic block on chromosome 10, containing seven genes, was found to be duplicated an average of 4.22 times in the KPGP donors. Included among these genes is *G protein regulated inducer of neurite outgrowth 2* (*GPRIN2*), which is associated with brain development and neurite outgrowth ^49^. Previous reports identified this duplication in Asian, European, and Yoruba populations (three-six copies), while no duplications were reported in the chimpanzee, orangutan, or gorilla ^22^. We also identified 444 CNVs conserved in 1000GP (Supplementary Table S10), which are probably shared East Asian CNVs and are not specific to Koreans. Five deletions and nine duplications were found to be enriched in the Korean population using the following criteria; i) odds ratio > 10 comparing with CNV ratio in any continents, ii) p-values < 0.01, and iii) more than five individuals in KoVariome. Phenotypic features were examined by searching genes against the OMIM database, resulting in the identification of three deletions and three duplications containing genes associated with known phenotypes (Fig. 5B). A high copy number deletion of *UDP glucuronosyltransferase family 2 member B17* (*UGT2B17*), which is associated with bone mineral density and osteoporosis^50^, was observed by comparing our Korean individuals with EUR, AFR, and AMR populations. This finding is consistent with previous studies which reported that 66.7% of Korean males have a deletion of this gene, compared to only 9.3% of Swedish males^51^. We also observed frequent deletions of *acyl-CoA thioesterase 1* (*ACOT1*), which functions to maintain the cellular levels of acyl-CoA and free fatty acids^52^. We identified the duplication of *hydroxycarboxylic acid receptor 2* (*HCAR2*) in 12% of the Koreans, which is associated with lipid-lowering effects^53^. We excluded the gene duplications of *NBPF15* and *HERC2* because they were located at the CNV break points. These CNVs will be useful for detecting Korean-specific genetic associations with specific phenotypes in future studies, which is especially important since CNVs are analyzed less often than SNVs even though they likely contain important disease-relevant variations.

**Figure 5.**
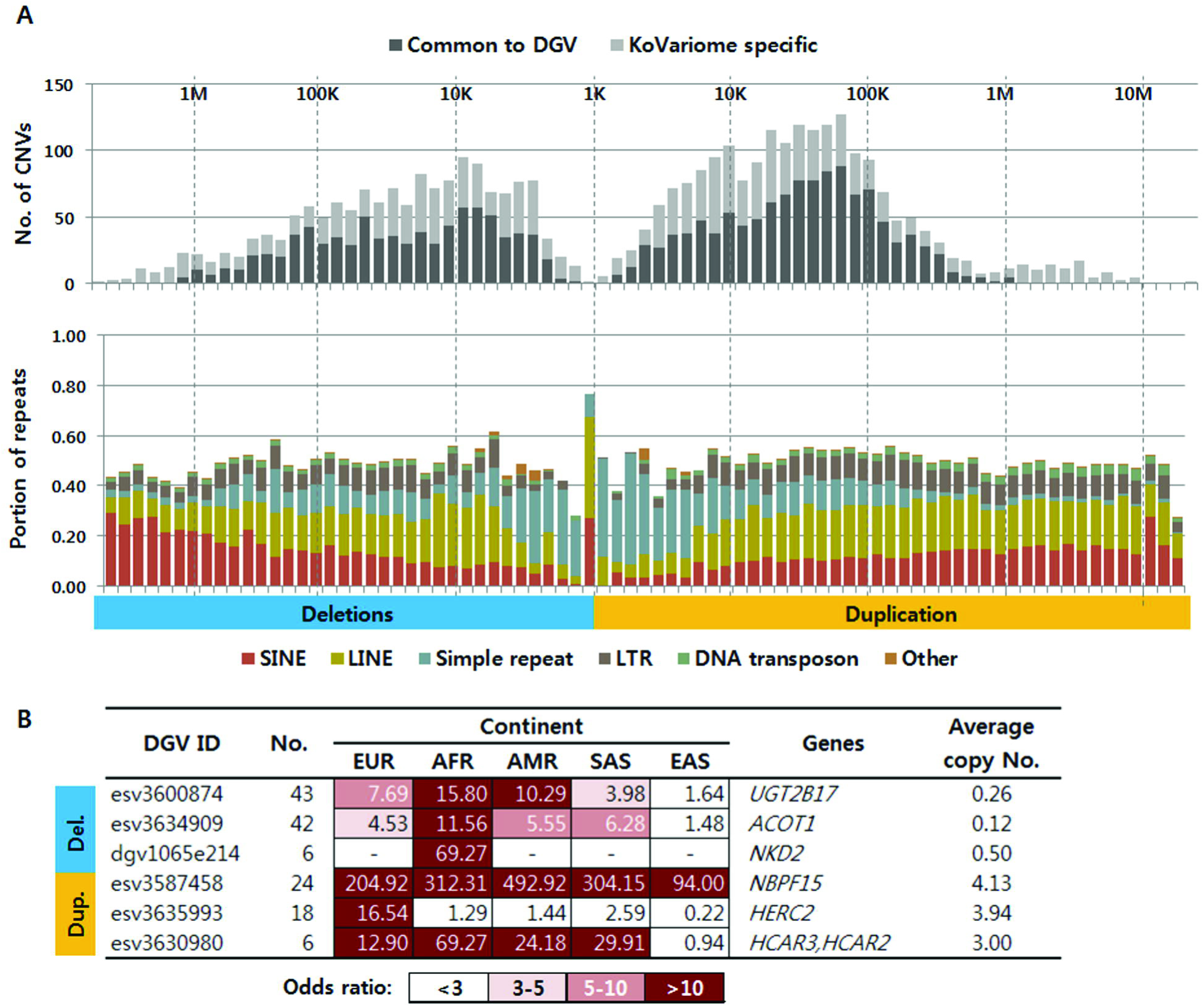
Properties of copy number variations in KoVariome. A. The number CNVs in the Korean population and the portion of the repeats in a specific length range. The conserved CNVs were defined by searching the Database of Genomic Variants (DGV) with 70% reciprocal overlaps. See the abbreviations of repeats in Fig. 4B. Korean enriched CNVs were identified by searching the CNVs reported in the 1000GP. No. represents the number of CNVs predicted in KoVariome. The heatmap represents the odds ratio of the CNVs compared to the CNV ratio in a specific 1000GP continental group. Associated genes were identified by searching the OMIM database. Abbreviations of continent group: European (EUR), African (AFR), Native American (AMR), South Asian (SAS), and East Asian (EAS).

**Table 5.** Copy number variations conserved in 50 Korean individuals.

| Chr. | Start | End | CNV<br>Types | Average<br>copy number | Genes <sup>a</sup> |
| --- | --- | --- | --- | --- | --- |
| chr2 | 132,964,050 | 133,121,849 | Dup. | 4.02 | *MIR663B*, *FAM201B*, *ZNF806*,<br>*ANKRD30BL* |
| chr10 | 46,222,900 | 46,946,499 | Del. | 1.0 | *PTPN20*, *FAM35BP*, *AGAP4*, *FRMPD2B*,<br>*FAM21C*, *BMS1P5* |
| chr10 | 46,946,200 | 47,150,299 | Dup. | 4.22 | *NPY4R*, *GPRIN2*, *CH17-360D5.1*,<br>*LINC00842*, *LOC102724593*,<br>*HNRNPA1P33*, *SYT15* |
| chr10 | 47,147,400 | 47,384,499 | Del. | 1.0 | *ANXA8*, *FAM35DP*, *LINC00842*,<br>*FAM25C*, *AGAP9*, *FAM25G*, *BMS1P6* |
| chr15 | 21,885,000 | 21,944,149 | Dup. | 6.4 | *LOC646214* |

<sup>a</sup> Genes in the identified CNV region. Chr. Chromosome; Dup. duplication; Del. deletions.

## Conclusions

To discover disease-causing genetic variants, researchers rely on comprehensive, population-specific databases containing the benign genetic variation present within specific ethnic groups. The KoVariome database was created to fill this need for the Korean population, and includes 5.5 TB of WGS data from 50 healthy, unrelated Korean individuals with corresponding health metadata. Using this database, we characterized all four variation types and identified 12.7M SNVs, 1.7M indels, 4K SVs, and 3.6K CNVs, many of which were novel or selectively enriched in the Korean population. Despite their close geographic proximity, the Korean population was shown to be genetically distinct from the Chinese and Japanese populations, highlighting the need for a Korean-specific variome to accurately identify rare disease variants in this population. Accordingly, KoVariome was used to predict candidate loci, inheritance patterns, and genetic risk for several diseases, including cancer, fibrodysplasia ossificans progressive, Van der Woude syndrome, type II diabetes mellitus, and spherocytosis. As this database grows and the accuracy of predicting disease associations increases, genetic tests will increasingly become increasingly routine components of precision healthcare. KoVariome will be an invaluable resource for biomedical researchers and health practitioners, and will directly benefit patients by ensuring they are presented with the most accurate genetic predictions of disease risks.

## Methods

### Sample collection and data distribution

Since 2010, the Korean variome data center (KOVAC) recruited volunteers for the Korea Personal Genome Project (KPGP). Consent was acquired from all participants in accordance with the Korean Life Ethics bill. In addition to providing a blood sample for WGS, each individual responded to a questionnaire regarding body characteristics, habits, response to 16 allergies, family histories, and physical condition related to 19 disease classes (Table S1). Genomic DNA was extracted using a QIAamp DNA Blood Mini Kit (Qiagen, CA, USA) and 69 WGS libraries were constructed using TruSeq DNA sample preparation kits (Illumina, CA, USA). Sequencing was performed using Illumina HiSeq sequencers following the manufacturer’s instruction. WGS data from 50 healthy unrelated Korean individuals were analyzed to create the KoVariome database, which was released through the national FTP portal server of the KOBIC (ftp://ftp.kobic.re.kr/pub/KPGP/) and distributed through GRF (http://pgi.re.kr) and Variome.net. All data analyzed in this study were deposited in NCBI SRA (PRJNA284338) and accessions for each sample were listed in Supplementary Table S2.

### Analysis of SNVs and indels

The WGS data were processed according to a protocol that was evaluated by the technical committee of the Korean Research Institute of Standards and Science (KRISS). Genomic resources were downloaded from UCSC Genome bioinformatics (http://hgdownload.cse.ucsc.edu/goldenPath/hg19/bigZips/), including the reference human genome (GRCH37/hg19), reference genes, and repeat annotations. Raw DNA reads were cleaned by Sickle (https://github.com/najoshi/sickle) with a quality score > 20 and read length > 50 bp. Cleaned paired-end reads were mapped to the human reference genome using BWA ^54^and indels were realigned and recalibrated after removing the PCR duplicates. Finally, we identified SNVs and indels for each individual using the GATK UnifiedGenotyper (ver. GATK-Lite-2.3-9)^55^. To improve the quality of identified SNVs, we applied SNV meeting criteria of: i) read depth (DP) is 20× or higher, ii) mapping rate is 90% or higher. Low-quality indels were removed from future analyses using the following criteria: i) quality score <27 and DP <6, ii) heterozygous indels with mapped allelic valance less than 0.3.

### Functional effect of the variants

To analyze the functional effects of variations, we implemented SnpEff-3.3^56^. The deleterious effects of the non-synonymous SNVs were obtained by searching dbNSFP (ver. 2.9.1), a portal database providing deleterious non-synonymous SNVs^57^. We then predicted the effects of each variant on protein function using SIFT, Polyphen2, PROVEAN, MetaSVM, and MetaLE, and further annotated variants using the Interpro_domain and COSMIC (Catalogue of Somatic Mutations in Cancer, ver. 71) databases. Previously reported SNVs and indels were identified using the dbSNP database (ver. 146). All variants shorter than 50 bp were then stored in this database^58^. The databases ClinVar (ver. 20161101)^59^ and OMIM (generated 2016-11-22)^60^ were searched to identify known pathogenic variants.

### Genetic distance calculation

The genetic distance (pi) between two samples was calculated using the following formula:

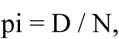

where D is the nucleotide difference between two samples and N is the number of compared positions. The sum of the nucleotide difference was calculated between two samples for each genomic position, which ranged from 0-1. A homozygous genotype composed of a reference allele was adopted as the genotype for uncalled sites.

### Multidimensional Scaling (MDS) analysis

Genotype data for 84 Chinese and 86 Japanese individuals were obtained from Phase 3 of the HapMap project^3^. A total of 1,387,956 SNV loci were merged with KoVariome. The PLINK program was used to remove the genomic loci with MAF < 0.05, call rates < 0.05, and SNPs in linkage disequilibrium blocks^61^. In total, 117,521 SNPs remained after filtering and were used in the MDS analysis. Five dimensional components were calculated in R with the distance matrix method “canberra” and MDS plots were generated using the MASS package ^62^.

### Accuracy of the SNVs

To measure the accuracy of SNV predictions, 35 individuals were genotyped with the Axiom^TM^ Genome-Wide East Asian (ASI) 1 Array (Affymetrix, Inc.). The accuracy and recalls were analyzed using a contingency table constructed with the presence and absence of the alternative alleles analyzed from our pipeline and the genotyping results from the Axiom^TM^ Genome-ASI 1 Array. The precision of calls was calculated by analyzing the concordance and denoted as true positive predictions (TP) from all predicted SNVs. The recalls were defined as TPs divided by the number of genotypes represented on the Axiom^TM^ Genome-ASI 1 Array. The genotype accuracies were measured by analyzing the concordance of the genotypes between the GATK prediction and the results from the Axiom^TM^ Genome-ASI 1 Array. The accuracy of the indel predictions were calculated by comparing genotypes between GATK predictions and the Axiom^TM^ Genome-ASI 1 Array.

### Structural variants

We applied two programs, BD^63^ and pindel^64^, to predict genome-wide SVs based on the discordant mate-pair and split-read information, respectively. From the bam files for each individual, insertions and deletions of a length between 100 and 1 Kb were predicted by pindel (ver. 0.2.4t) and those longer than 1Kb were predicted by BD (ver. 1.4.5)^65^. We next constructed unassembled genomic blocks (‘N’) from the hg19 reference genome and examined the SVs that overlapped with these unassembled genomic regions. From this analysis, we discovered a high portion of spurious SVs in these regions (Supplementary Fig. S5), with the majority of them >100M in size. The following criteria were used to filter out spurious SVs; i) reciprocally > 10% overlaps between SVs and un-assembled genomic blocks, ii) ‘N’s more than 50% coverage of SVs, and iii) more than 2 un-assembled genomic blocks in the predicted SVs. After filtering, we clustered SVs that reciprocally overlapped > 70% in any individual. Unified SVs were defined by the average start and end positions in each SV cluster. The novelty of each SV was defined by comparing unified SVs with those in the DGV^66^, with 70% reciprocal overlaps.

### Copy number variations

CNVs were predicted with FREEC (ver. 10.6) using window size =100, step size =50, and breakpoint =0.6 ^48^. The spurious CNVs were enriched in >1M in length (Figure S7), which were filtered using the same criteria described in the SV methods above. Unified CNVs were constructed by merging individual’s CNVs that reciprocally overlapped by >=70%. The start and end positions of the unified CNVs were defined as average position of the original calls. Known CNVs were defined by comparing with CNVs in the DGV database^66^.

## Additional information

### Data resource access

http://variome.net, http://kpgp.kr, http://koreangenome.org SNP data have been deposited in dbSNP under batch_id 1062763.

## Acknowledgements

The authors thank many people not listed as authors who provided analyses, data, feedback, samples, and encouragement. Especially, thanks for Sunghoon Lee, Taehyung Kim, Sanghoon Song, Sangsoo Kim, and George Church. This work was supported by the 2014 Research Fund (1.140113.01 and 1.150014.01) and the 2016 Research Fund (1.160052.01) of Ulsan National Institute of Science & Technology (UNIST) and supported by the Ministry of Trade, Industry & Energy (MOTIE, Korea) under Industrial Technology Innovation Programs ‘National Center for standard Reference Data’, No.10075262. This work was also supported by the Research Fund (14-BR-SS-03) of Civil-Military Technology Cooperation Program and the Next-Generation Information Computing Development Program through the National Research Foundation of Korea (NRF) funded by the Ministry of Science, ICT & Future Planning (NRF-2016M3C4A7952635).

## Author Contributions

J.B. and B.K. planned and designed this study. J.K., S.J., J.J., H.-M.K., H.K., and O.C. processed the sequencing data. Y.K, J.B., and J.J. contributed to recruitment of individual for whole genome sequencing. J.K., S.L., J.B., Y.C., and J.W., contributed to the interpretation of the data. J.W., C.K., H.L., B.K., K.H., I.K., J.E., J.B., K.C., and J.E. wrote and reviewed draft. All authors commented on the manuscript and approved the final version to be submitted.

## Supplementary information

1. Supplementary Tables
2. SupplementaryFigures

## Comments

By submitting a comment you agree to abide by our Terms and Community Guidelines. If you find something abusive or that does not comply with our terms or guidelines please flag it as inappropriate.

